# Spontaneous Phase Separation of Cocultured Cell Mixtures *In vitro*

**DOI:** 10.1101/152819

**Authors:** Sebastian V. Hadjiantoniou, Maxime Leblanc-Latour, Maxime Ignacio, Cory S. Lefevbre, Gary W. Slater, Andrew E. Pelling

**Affiliations:** Department of Biology, Gendron Hall, 30 Marie Curie, University of Ottawa*, Ottawa, ON, K1N 6N5,* Canada; Department of Physics, 598 King Edward, University of Ottawa*, Ottawa, ON, K1N 6N5,* Canada; Institute for Science, Society and Policy*, Desmarais Building, 55 Laurier Ave. East,* University of Ottawa*, Ottawa, ON, K1N 6N5,* Canada

**Keywords:** Biophysics, Cell biology, Molecular Biology, Coculture, Adhesion

## Abstract

During Embryogenesis, cells undergo constant organizational remodelling. Biochemical and biophysical guidance cues act in tandem to guide migration and morphogenesis into distinct cellular patterns. It has been shown that various cell types will express different configurations of cellular adhesion molecules known as cadherins and integrins. Cocultured *in vitro* experiments have focused on revealing the extensive genetic expression profiles that modulate embryogenesis whilst overlooking the physical cell-cell and cell-substrate interactions that influence organization. We demonstrate that NIH3T3 and MDCK cells undergo a spontaneous phase separation when cocultured *in vitro* and that this phenomenon occurs through purely physical binding energies. A Monte Carlo simulation model of a mixture of cells with different cell-cell and cell-substrate binding energies reveals that the spontaneous phase separation occurs due to the minimization of interfacial free energy within the system. Cell-cell and cell-substrate binding plays a critical role in cell organization and is capable of phase separating different populations of cells in vitro.

## Introduction

At the heart of embryogenesis lies an extensively complex list of cues which drive the cohesive formation of organized tissue^1^. Efforts to elucidate these governing mechanisms have resulted in identifying two key proponents in development: differentiation and cellular organization. Differentiation is ultimately controlled through highly regulated transcriptional activity^2^. On the other hand, explaining the mechanisms of cellular organization has required the examination of the mechanisms that impact cell-cell and cell-substrate interactions. It is for this reason that Steinberg et al. ^3–5^ proposed the differential adhesion hypothesis (DAH), which suggests that morphogenetic sorting forms through tissue interfacial free energies arising from cellular adhesive interactions ^6^. In other words, cells possess cellular adhesion molecules (CAMs) known as integrins and cadherins, responsible for cell-substrate and cell-cell binding, respectively. Depending on the cell type and cellular event, different configurations of CAMs will be present on the surface of the cell. These configurations lead to relative intensities of intercellular adhesion, which serve as a set of morphological determinants creating highly organized cellular patterns through passive or active motility. The physical phenomenon of adopting the lowest free energy configuration was originally shown in a study whereby amphibian embryonic cells were disassociated, mixed, and subsequently allowed to re-associate. The outcome showed a spherical reformation with cells that re-aggregated into their embryonic position^4^. More recent studies have shown similar examples of the influential effects of differential cell to cell vs cell to substrate adhesion. Mouse embryonic stem cells were seeded into channelled topographies whereby after 48hrs, the cells demonstrated preferential cell-cell adhesion and formed spherical embryoid bodies rather than flat island-like aggregates commonly seen in flat two dimensional culture vessels^7^.

Despite the clear multicellular environment of native tissue, co-cultured systems, whereby multiple cell types are cultured together in vitro, have primarily focused on analyzing changes in gene expression ^8–11^. For example, some progress has been made at characterising the relationship between carcinoma and stromal cells as cell-cell communication has been considered to play an important role in triggering cancerogenesis^12–14^. Many of these coculture systems however prefer a paracrine-only interaction by utilizing separated adherable membranes ^8^. Although this method elucidates potential biochemical influences, it prevents any physical cell-cell interaction from occurring. As such, cells may display patterned growth that would be solely attributed to reaction-diffusion mechanisms while completely ignoring the effects of differential adhesion. Other studies have explored differential adhesion via synthetic biology strategies. By utilizing the L-929 murine fibroblast cell line which expresses no known cadherins and displays no aggregation when placed in hanging drops, Steinberg et al. were able to modulate exogenous cadherin expression to induce predictable patterned formations^15^. Similar results were obtained by Cachat et al., whereby aggregated spheroids displayed cell partitioning (termed phase separation) depending on cadherin expression patterns^16^. Despite the extensive work via synthetic biological approaches to demonstrating cadherin based differential adhesion, utilizing endogenously expressed cadherin based sorting has yet to be demonstrated.

In this communication, we use unaltered cell lines in coculture and demonstrate spontaneous phase separation in mixtures of two distinct cell types on flat two dimensional topographies. To achieve this, we specifically selected two distinct cell types with different endogenously expressed cadherin profiles. NIH3T3 fibroblasts are highly motile, transiently adherent cells whilst MDCK epithelial cells form strong cell-cell tight junction bonds. In vitro, these cocultured cells quickly underwent a phase separation. MDCK cells randomly dispersed, eventually forming a monolayer on the substrate whilst NIH3T3s slowly aggregated into patterned formations. We also illustrate how selectively inhibiting signalling cascades that modulate binding affinities between cells and substrate resulted in statistically different cellular patterning formations. Furthermore, to validate that the observed phenomenon could in fact be achieved through purely physically driven forces, we developed a kinetic Monte Carlo computer simulation model of our experimental system. By mapping cell-cell and cell substrate adhesion into binding energies and minimizing the interfacial free energies between cells, our model was able to replicate our in vitro coculture results. Taken together, our study suggests how highly influential differential adhesion is in mixtures of cells in-vitro. Moreover, even when key biological mechanisms that dictate the morphology and sensory perception of the cell are inhibited, fundamental biophysical interactions are capable of driving cellular phase separation.

## 2. Materials and Methods

### 2.1 Cell culture and drug studies

NIH3T3 mouse fibroblast cells (ATCC® CRL-1658™) and Madin Darby Canine Kidney (MDCK) epithelial cells were cultured in high glucose DMEM containing 10% Fetal Bovine Serum (FBS) and 1% penicillin/streptomycin antibiotics (all from Hyclone). The cells were cultured at 37 °C and 5% CO2 in 100 mm dishes. Monoculture experiments were performed by seeding ~20 000 cells/cm^2^ on tissue culture treated petri dishes for 48 hrs. Cocultures of NIH3T3 and MDCK cells were thoroughly mixed and seeded at equal densities of ~10 000 cells/cm^2^ and subsequently imaged in the same manner as monoculture experiments. Inhibition studies of ROCK (Y-27632; 10 μM in dH_2_O, Sigma, Catalogue #Y0503), Myosin II (Blebbistatin; 10 μM in DMSO, Sigma, Catalogue #B0560) and mDia (SMIFH2; 10 μM in DMSO, Sigma, Catalogue #S4826) were all performed by exposing both cell types for the 48 hrs incubation time period.

### 2.3. Immunofluorescence staining and microscopy

Coculture experiments were performed by pre-loading NIH3T3s and MDCKs with CellTracker Green CMTPX dye (Invitrogen, Catalogue #C34552) and CellTracker Red CMFDA dye (Invitrogen, Catalogue #C7025), respectively. After a 48 hr incubation period, the dyes were removed and the cells were rinsed three times with DMEM. Cells were then trypsinized and plated on the microchanneled grooves as describe in section 2.2. After 48 hrs, cells were fixed with 3.5% paraformaldehyde and stained with Dapi (Invitrogen, Catalogue #D1306).

Monoculture experiments were fixed after 48 hrs incubation with 3.5% paraformaldehyde and permeabilized with Triton X-100 at 37°C. Cells were stained for actin using phalloidin Alexa Fluor 488 & 546 (Invitrogen, Catalogue # A12379 & #A22283) and DNA was stained using DAPI. A full protocol has been published previously ^17^. Samples were then mounted using Vectashield (Vector Labs) and a #1 coverslip placed on top. Samples were then inverted and imaged with a Nikon Ti-E A1-R high-speed resonant laser scanning confocal microscope (LSCM) with a phase contrast 10x NA0.3 objective or a DIC 60x NA1.2 water immersion objective. For high resolution three dimensional image capture of cocultured cells, NIH3T3 GFP (Cedarlane, #AKR-214) were used with MDCKs and subsequently fixed, stained and imaged as described above.

### 2.4 Parallel Plate Flow Assay

A polydimethylsiloxane (PDMS) microfluidic flow chamber, 1000 mm in width, 160 μm in height and 20 mm long, was produced using standard microfabrication techniques. Liquid-state 1:10 ratio PDMS was poured on the master and cured for 2 hours at 70°C. Inlets and outlets, 0.75 mm in diameter, were punched at both ends of the chamber. The PDMS layer was bonded to a standard glass microscope slide after being plasma treatment (50 W for 30 sec), creating an enclosed channel. The entire chamber was submerged in media and cells were then pipetted at a seeding density of ~4 million/ml each. Cells were cultured inside the chamber overnight. Y-27632 (10 μM) was added to the cells prior to seeding so as to expose the cells until the flow assay. The cells were exposed to an average flow of 1μl/s, for 30 min, which corresponded to a shear stress at the wall of the chamber of 2 Dyne/cm^2^ [42]. The flow rate was then increased to 13 μl/s (30 Dyne/cm^2^) for an additional 15 min. Images were captured every 5 minutes from the same region of interest before, during and after exposure to shear stress.

### 2.5 Image and statistical analysis

A Kolmogorov-Smirnoff (K-S) test was performed for each study case in order to determine if they exhibited patterned formation. Analysis consisted of partitioning the image samples into 100 quadrats, whereby the number of each cell type per quadrat was recorded and the corresponding cumulative distribution function was calculated^18^. The absolute maximum difference D compares *F_exp_* and the cumulative distribution function of a Poisson distribution (*F_Pois_*.):

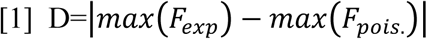

Each D was computed independently for epithelial cells (type 1) and fibroblast cells (type 2) in the *in vitro* experiments and simulation model. For a value of D above 0.195, we rejected the null hypothesis of a random distribution at a 99.9% confidence threshold, and concluded that the cells exhibit a patterning. For a D value below 0.136 (corresponding to the 95% confidence threshold), we concluded that the cells were randomly distributed. All statistical analyses were carried out using *ImageJ*. In addition to the K-S distribution test, we quantified the phase separation of the two cell types in the computational simulation.

### 2.6. Kinetic Lattice Monte Carlo Simulations

Through the use of Kinetic lattice Monte Carlo (KMC) simulations^19,20^, we have modeled the morphological “evolution” of a binary system with *N* = *N*_1_ + *N*_2_ cells initially deposited on a two dimensional (2D) flat substrate, where the N_1_ and N_2_ type cells are analogous to NI3T3s and MDCKs, respectively, used in both the untreated and Y-27632 *in vitro* experiments. Each cell occupies a three dimensional (3D) cubic lattice site of volume *α*^3^ = 1. Initially, each lattice site on the surface has a 0.95 uniform probability of being occupied by a cell of type 1 or type 2 (0.475 each) and a 0.05 probability of being unoccupied (in our case, an unoccupied lattice space is analogous to a cell media element). We used a simulation box of volume *V* = *L_X_* · *L_Y_* · *L_Z_* (50·50·10) with lateral periodic boundary conditions in x and y. For boundary conditions in z, the position z = 0 represents the flat substrate whilst a sufficiently large L_z_ was chosen to ensure that no cell touch the upper boundary condition during the simulation.

Cells may only move to a first nearest-neighbor free lattice site^21^ (corresponding to a lattice site being occupied by cell media), such that two cells cannot simultaneously occupy the same site. However, cell motion can only be accepted if the moving cell remains in the vicinity of either another cell or the substrate (vicinity is defined as being a first- or a second -nearest neighbor); in other words, cells are not allowed to swim freely in solution.

Cells of type i, where *i*=1,2, can form bonds with cells of the same type (homotypic binding energy: *ε_ii_*) or of the other type (heterotypic bond energy: *ε_if_* with *i*≠*j*). We also account for the interactions between the cells and the substrate (substrate binding energies: *ε_is_*,) and we set the interaction energy with the surrounding media at zero^22^. Moreover, one does not treat the substrate energy since it has no internal degree of freedom. For each cell *i*, we define an energy barrier to be overcome to complete a move, *E_b_* = *n_i_*_1_*ε_i_*_1_ + *n_i_*_2_*ε_i_*_2_ + *n_is_ε_js_*, where *n_ij_* (j = 1,2) are the number of first neighbors of each type. Given this energy barrier, each cell is assigned a transition rate given by

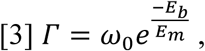

which only depends on the initial energy configuration, a constant attempt frequency ω_0_ and the typical energy of biological fluctuations *E_m_* ^23^. The latter characterizes cell motility driven by cytoskeleton motion (*E_m_* is equivalent, from a thermodynamic perspective, to the thermal energy *K_b_T*). As defined by [3], the transition rates allow to reach the equilibrium state which corresponds to a configuration with a minimum surface/interface energy^24,25^.

### KMC parameters

All the energy parameters are expressed in *E_m_* units. The values of the binding energies were chosen in order to semi-quantitatively reproduce the experimental observations, in both untreated and Y-27632 experiments. We assumed that the binding energies between homotypic cells are comparable to *E_m_*. MDCK cells (type 2) are known to strongly interact with the substrate and form a 2D monolayer. Based on this, the binding energies have to be chosen in order to verify 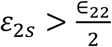 and *ε*_2*s*_ > *E_m_*, *ε*_2*s*_ ≫ *E_B_*. On the other hand, NIH3T3 cells (type 1) can form homotypic 3D clusters on the substrate and over the MDCKs. This condition suggests that the binding energies have to be chosen to verify that 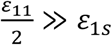, 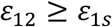 and 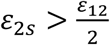. To simulate the effect of the Rock inhibition, we used the untreated parameters as a set baseline and lowered *ε*_1*s*_ to replicate the inhibitory effects of Y-27632. Finally, to be consistent with the DAH whereas phase separation is expected between cell type with non-equal binding energies, we must verify that 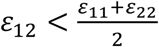. Numerical values are shown in Table 1.

**Table 1:** Parameter values for the binding energies. Subscript i refers to cell type 1 and 2, s to a substrate lattice site”

| Binding energy | Values<br>(in units of $E_m$)<br>Untreated / Y-27632 |
| --- | --- |
| $\epsilon_{11}$ | 1.05 / 1.05 |
| $\epsilon_{22}$ | 4 / 4 |
| $\epsilon_{12}$ | 1.5 / 1.5 |
| $\epsilon_{1s}$ | 0.5 / 0.3 |
| $\epsilon_{2s}$ | 5 / 5 |

## 3. Results

### 3.1 The role of the cytoskeleton in cocultured phase separation

To explore the behavioural interaction between cells, NIH3T3 and MDCK cells were co - cultured for 48 hrs. NIH3T3 fibroblasts and MDCKs epithelial cells were preloaded with a green and red fluorescent cell tracker, respectively, permitting visual differentiation during imaging. As can be seen in Fig 1A, there is a clear phase separation of fibroblast cells (green) compared to the epithelial cells (red). To assess the involvement of key cytoskeletal proteins, we systematically performed the same set of experiments with selective inhibitory drugs. At a concentration of 10 μM, Y-27632, Blebbistatin and SMIFH2 were each added to their respective samples seeded with ~25 000 cell/cm^2^ for 48 hrs. As shown in Fig 1A, inhibition of actin contractility (Blebbistatin) and actin polymerization (SMIFH2) had no effect on the behavioural interaction of both cell types in coculture. Inhibition of cytoskeletal signalling cascades (Y-27632) however presented more definable changes in patterns of cellular formation. Whilst MDCK epithelial growth remained widely dispersed in the form of a monolayer, NIH3T3s aggregated into larger islands connected via strands of cells. Further LSCM analysis reveals that there is in fact a monolayer of MDCK cells below the patterned NIH3T3s (Fig 1b). These results are in contrast to NIH3T3 and MDCK cells cultured in isolation in which they formed a mesh-like network or a monolayer, respectively, as has been long established and observed^26^ (Supp Fig 1).

**FIG. 1.**
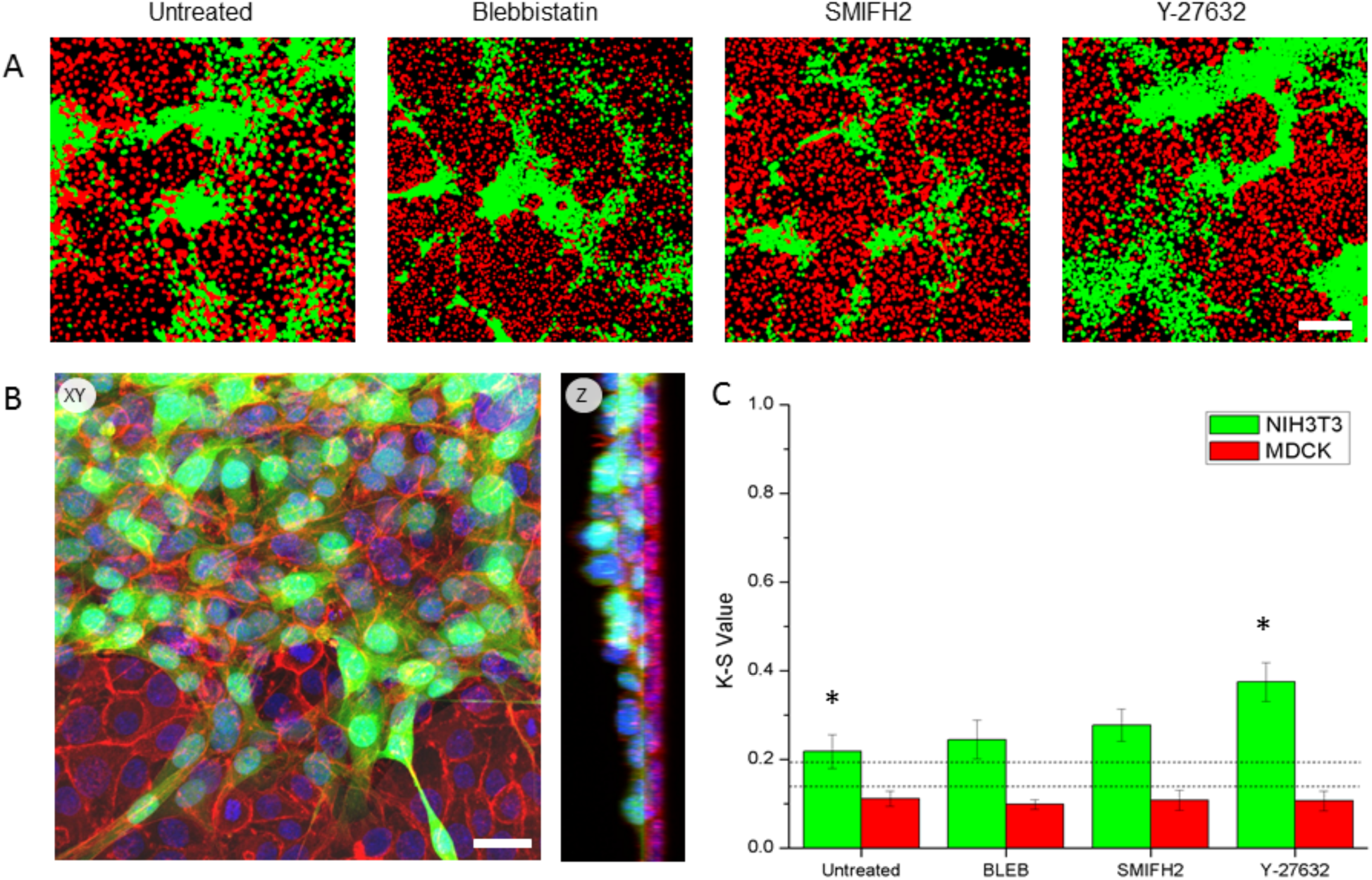
Co-cultured NIH3T3 and MDCK on flat substrates. (A) NIH3T3 (green) and MDCKs(red) co-cultured and seeded onto a flat substrates for 48 hrs. Images have been thresholded to amplify the contrast between cell types. Distinct patterns of cell phase separations are apparent in all conditions. (B) Kolmogorov-Smirnoff test analyzing the presence of patterned behavior in both cell types. The lower (0.136) and higher (0.195) critical values (dotted lines) correspond to the confidence threshold for which the null hypothesis of a random distribution can be rejected. Therefore, above indicates patterned formation whilst below suggests random cell dispersion. In all cases, NIH3T3s display patterned cell behavior whilst MDCKs present a random cellular distribution pattern. Statistically significant (p<0.05) results were found when comparing untreated cells to Rock inhibited (Y-27632) patterning. Scale bar = 200μm (For interpretation of the references to color in this figure legend, the reader is referred to the web version of this article). (C) Orthogonal view of NIH3T3 (green) and MDCK (red) co-cultured on a flat substrate. Z projection rendering of confocal stacks reveals the layered patterning as NIH3T3s directly on a monolayer sheet of MDCK cells. (Scale bar = 25μm)

In order to quantify and assess this visual cell patterning (Fig 1c), a Kolmogorov-Smirnov (K-S) test was used. This includes performing a quadrat analysis to compare the cell frequency distribution to a random Poisson distribution by measuring the absolute, largest difference between cumulative frequencies, designated as the D statistic-value which has been described in detail previously^16^. Briefly, the D statistic-value is a dimensionless parameter that allows one to quantitatively determine if a cell population is randomly distributed (D < 0.136) or not (D > 0.195). In coculture, the D statistic value for MDCKs was 0.112±0.02 (p>0.05, n=7), indicating that the distribution is not statistically different from a random spatial distribution. As expected, NIH3T3s cells have a much higher D value of 0.218±0.04 (p<0.05, n=7), confirming that their distribution is patterned and suggesting that there is a governing mechanism controlling their aggregation.

Quadrat analysis indicates a significant patterning difference in NIH3T3s with a D statistic value of 0.37±0.04 (p<0.01, n=8) when cultured with Y-27632 compared to the untreated. Despite the observed changes in patterning with Y-27632, NIH3T3 cells demonstrated only minor variation in their patterning behaviour with D values of 0.244±0.04 (p>0.05, n=8) and 0.277±0.04 (p>0.05, n=9) for Blebbistatin and SMIFH2, respectively. MDCKs however demonstrate relatively no patterning change among any of the selective inhibitions with a consistent D value of 0.9-1.1 (Fig 1c).

### 3.2 Cell substrate binding affinity in cocultured cells

Based on these results, we hypothesized that Y-27632 treated cells underwent inhibition of cell-substrate adhesion leading to increased aggregation and phase separation. To test this hypothesis, we performed a flow assay^27^ with a cocultured system confined within a microchannel (Fig 2 a,b). At a shear force of 2 dynes/cm^2^, untreated NIH3T3s adhered significantly more than Y-27632 treated cells with 58% of cells remaining compared to only 29% (p<0.05, n = 3) after 30 minutes. When increasing the flow to a maximum shear force of 30 dynes/cm^2^, 40% of untreated cells remained attached compare to 23% after treatment with Y-27632. It is important to note that under all conditions MDCK cells remained unaffected. This result further confirms that the cell-substrate binding affinity of MDCKs is significantly higher than that of NIH3T3s.

**FIG. 2.**
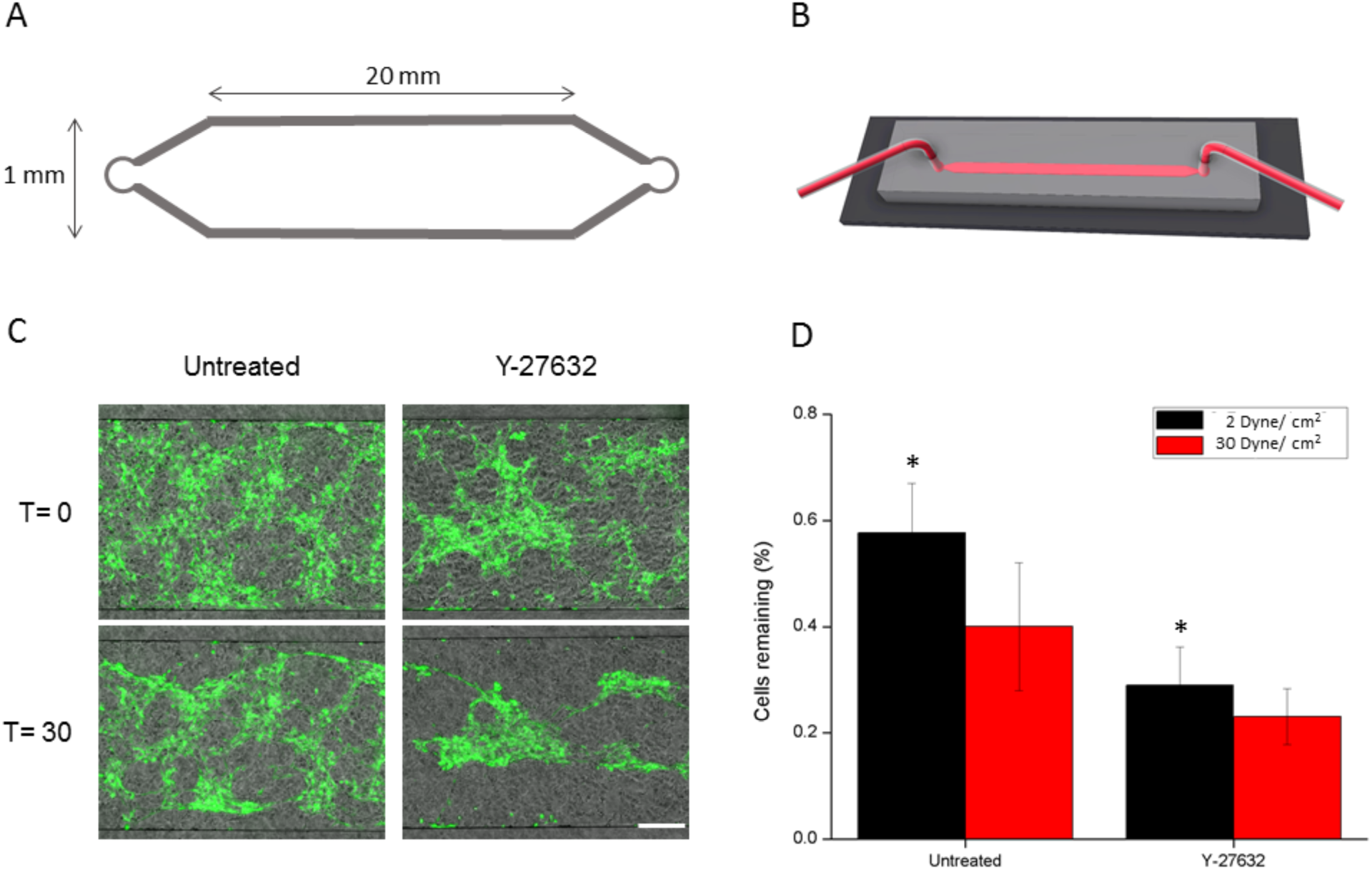
Flow adhesion assay of cocultured NIH3T3s and MDCKs. (A) Flow chamber design dimensions, L= 20mm, W= 1mm and H=160μm. (B) Graphical rendering of PDMS mold inverted and bound onto a glass slide with microfluidic tubing inserted into the 0.75mm inlets. Typical adhesion assay where cells were exposed to shear stress over a period of 30 min. Cells were counted before and after flow to quantify changes in cell-substrate adhesion strength (scale bar = 250 μm). (A) NIH3T3s (green) and MDCKs (phase) before (T=0) and after (t=30) 1μl/s of flow (2 dynes/cm2). (B) Ratio of cells remaining on the glass substrate after exposing to a shear stress of 2 & 30 Dyne/cm2 over a period of 30 min. There is a significant difference between the untreated and Y27632 treated cells (p<0.05).

### 3.3 Simulating Patterned Adhesion

It has been well established that different cell types express their own combinations of cell adhesion molecules (CAMs)^28,29^. Through specific integrin isoforms and cadherin binding, differential adhesion energies can form complex cell patterning^30–32^. To assess whether the experimentally observed phenomenon of cell phase separation between NIH3T3s and MDCKs is achievable through differential adhesions alone, we developed a computer simulation model. Our Kinetic Monte Carlo simulation utilizes properly chosen binding energies between cells (*ε*_11_, *ε*_12_, *ε*_22_) and substrate (*ε*_1*S*_, *ε*_2*S*_) to model the various interactions taking place during cell migration (Table 1). This simulation is not meant to fully represent the biological complexities present in vitro, but to ascertain whether the phase separation observed in vitro can be obtained through purely physical interactions. When simulating co-cultures on flat two dimensional topographies the results display the same phase separation phenomenon to cells cultured in vitro (Fig 3A). Simulated co-cultures with the chosen NIH3T3 (ε_11_=1.05, ε_1s_=0.5) and MDCK (ε_22_=4, ε_2s_=5), (ε_12_=1.5) energy parameters were found to replicate the in vitro observations. NIH3T3 patterning was statistically significant with a K-S D value of 0.26±0.01 (p<0.05, n=5) (Fig3B). Simulated MDCKs also showed identical cell dispersion to that observed in vitro, with a D value of 0.10±0.02 (p>0.05, n=5).

**FIG. 3.**
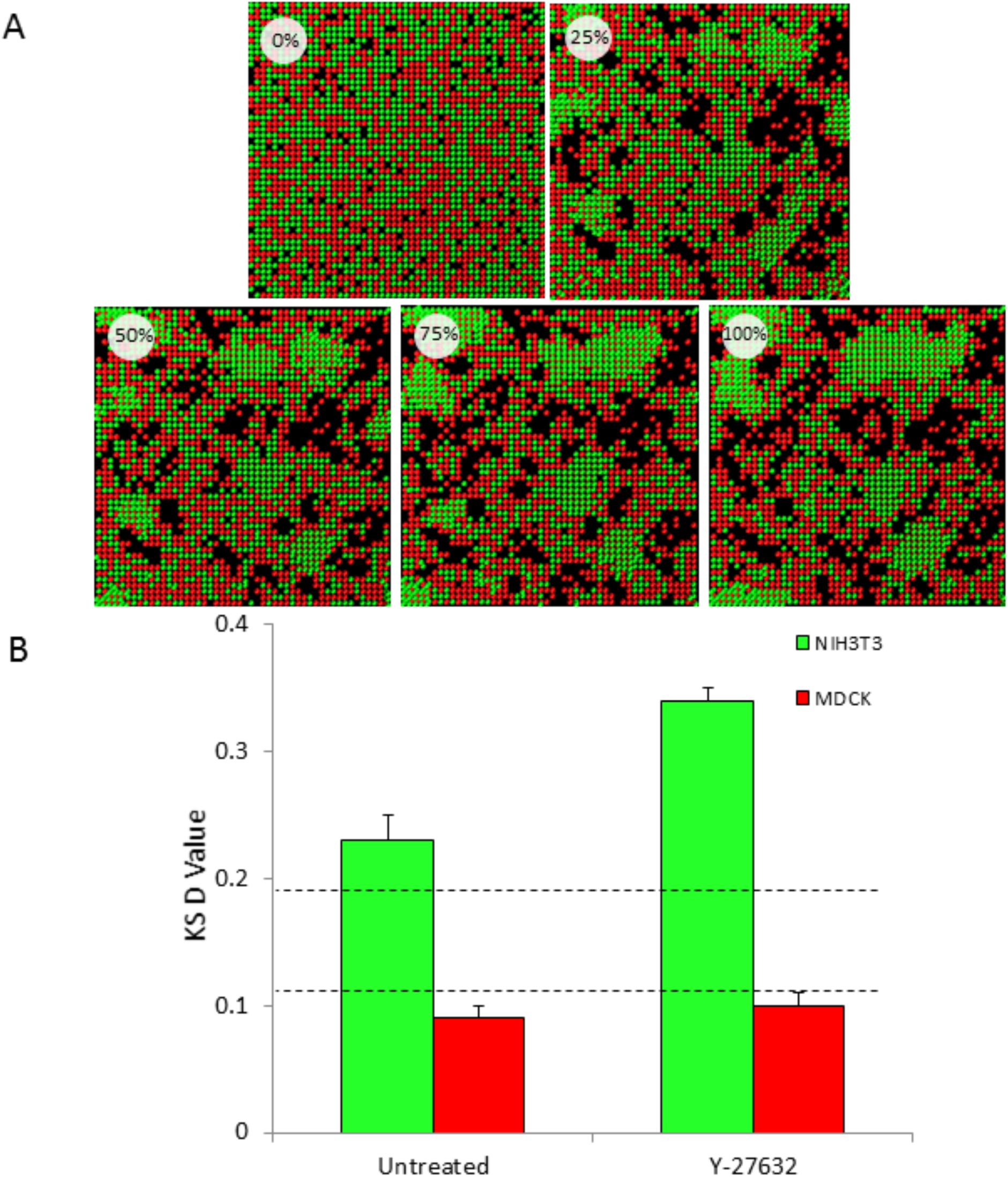
Simulation of co-cultured cells utilizing the differential adhesion model of patterning. A kinetic Monte Carlo simulation whereby cells undergo stochastic diffusion and interact with one another based on their binding energy parameters. These parameters (*ε*_11_, *ε*_22_, *ε*_12_, *ε*_1s_, *ε*_2s_) represent the cell-cell and cell-substrate adhesion of NIH3T3s/MDCK and have been modeled accordingly. (A) Timelapse sequence of rendered simulation for NIH3T3s (green) and MDCKs(red) co-cultured onto a flat substrate. Similarly, to experimental results, the differences in adhesion energies between cell types create a two layered cell phase separation whereby MDCKs are widely dispersed on the bottom whilst 3T3s display aggregated islands on top. (B) Kolmogorov-Smirnoff test analyzing the presence of patterned behavior in both cell types. The lower (0.136) and higher (0.195) critical values (dotted lines) correspond to the confidence threshold for which the null hypothesis of a random distribution can be rejected. Therefore, above indicates patterned formation whilst below suggests random cell dispersion. Simulated NIH3T3s on two dimensional substrates display patterned cell behavior whilst MDCKs present a random cellular distribution pattern. (For interpretation of the references to color in this figure legend, the reader is referred to the web version of this article).

After identifying the binding parameters that replicate in vitro observations, we modulated them in accordance to the known biological effect of Y-27632. By merely decreasing the binding energy of NIH3T3 substrate adhesion (ε_1s_) from 0.5 to 0.3, we observed the same K-S D value of 0.33 ± 0.02 (p<0.05, n=5) as in vitro experiments. Many of the key observations made in vitro were also observed in our simulations. For instance, MDCKs adhere and form strong cell-substrate and cell-cell bonds whilst NIH3T3s migrate until they have contacted an adjacent cell of the same type. Over time, they form large island like aggregates (25μm high, corresponding to ~2-3 cells) positioned above the monolayered MDCK cells (Fig 3a). To compensate for the simulation model not taking into account cell duplication, we performed the same set of experiments with the addition of thymidine (2mM). As a deoxynucleoside, thymidine causes cell cycle arrest and is often utilized for cell synchronization^33^. Supplementary Fig 2a demonstrates that co-cultured cells under cell cycle arrest display the same patterning effect shown in mitotically active cells. KS test (Supplementary Fig 2b) quantitatively shows a statistically significant pattern in NIH3T3 cells (D value = 0.345±0.04, P<0.05, N= 3) whilst MDCK cells remain randomly dispersed (D value = 0.11±0.02, P> 0.05, N=3).

## Discussion

Although identified as the inherent governing mechanisms for cell patterning^5,31,34–36^, the differential adhesion and the morphogen gradient models remain experimentally bifurcated. Experiments focusing on cocultured systems have rarely permitted cells to not only biologically interact but to physically interact as well. In this study, we utilize a simple co-culturing strategy with cell types expressing different cadherin subtypes to examine cell patterning formation.

To investigate this phenomenon, we cocultured NIH3T3 fibroblasts and MDCK epithelial cells onto two dimensional flat topographies. After 48hrs incubation, the initial melange of cells had undergone a distinct cell phase separation creating a two-layered system. The first layer was a well dispersed epithelial monolayer of MDCKs. The second layer, on the other hand, was a complex structure of aggregated NIH3T3s throughout the system. As can be seen in Fig 1a, this aggregating behaviour is unlike their normal growth pattern (Supplementary Fig 1b), which usually forms large meshes consistent with connective tissue. The morphology and behaviour of MDCK cells however appears unaffected in coculture. To further simplify the patterning system from the dynamic biological complexities presented by cell proliferation, we plated a high density 1:1 mixture of cells types with the addition of a mitosis inhibitor. Mitosis inhibited cocultures (Supplementary Fig. 2) demonstrate the exact same cell patterning which suggests that this phase separation does not evolve from cell division. To further investigate the potential influence of biological mechanisms in this phase separation phenomenon, we performed a set of cytoskeleton inhibition experiments. Preventing myosin II contraction which has been shown to be a key protein in migration and tissue architecture demonstrated a slight change in altered patterning behaviour however this was not statistically significant (p>0.05). The same is true for when inhibiting de novo forming actin polymerization. Two hypotheses can be inferred from these results. First, the spontaneous patterning observed in coculture is not actively guided via migration, as this would require a properly functioning cytoskeleton. Studies have shown how exposure to Blebbistatin and SMIFH2 causes substantially decreased migration in addition to lowering focal adhesion turnover^37–39^. Secondly, although inhibition of actin contraction and polymerization did occur through myosin II and formin, compensatory mechanisms mitigated the overall effects on cell organization. Cross-talk between effector pathways and feedback inhibition is ubiquitous in normal signal transduction. When signaling is blocked, regulatory loops can up-regulate parallel pathways to compensate, permitting the cellular response to be dynamic and adaptive^40^. It is for this reason that the statistically different results observed with Y-27632 inhibition are highly revealing of the mechanisms at play. Rock is a high level pleotropic regulator of multiple signalling pathways mediating cytoskeleton reorganization^41^, stress fiber formation^42^, cell contraction and cell polarization^43^. Its inhibition results in a widespread downregulation of the various direct and indirect pathways regulating the cytoskeleton^43^. The increase in cell patterning suggests that cytoskeletal involvement is unlikely and that some other governing mechanism is driving the observed effect.

It has been established that Rock is involved in the maturation of focal adhesions (FA). These FA are multiprotein complexes serving as transmembrane links between the extracellular matrix and the actin cytoskeleton. During the initial stages of adhesion, FA maturation involves a sequential cascade of compositional changes^44^. Rock, on the other hand, promotes FA maturation through tension induced conformational changes^45^. This conformation change allows the recruitment of key proteins including talin, paxillin and vinculin into the FA complex. As a result, the cell is capable of forming mature adhesion sites strengthening their cell-substrate binding affinity. Inhibiting Rock via Y-27632 may sufficiently reduce cell-substrate affinity in NIH3T3s, causing cells to depend more highly on their cell-cell binding properties for adhesion. This is supported by our computer simulation results which showed that the impact of Y-27632 can be reproduced simply by lowering the cell-substrate adhesion from 0.5 to 0.3: modulating this binding adhesion induced an increase in the phase separation of cocultured cells as observed in vitro.

Rock has also been shown to be intimately involved in the regulation of cadherin ligation in epithelial cells^46^. As an epithelial cell line, MDCKs possess the E-cadherin superfamily of cellular adhesion molecules. These transmembrane glycoproteins mediate specific cell-cell adhesion, functioning as key molecules in the morphogenesis of a variety of organs^47,48^. Ecadherins are not passively distributed on the surface of the plasma membrane, but rather localize at cell-cell contacts in response to adhesion. This focalizing mechanism is dependent on functionally linked cytoplasmic regulators, of which Rock is involved^46^. During adhesion, two distinct patterns of cadherin clusters form, fine punctate and larger streak-like macroclusters. Inhibition of rock activity preferentially abolishes the macroclusters which form the termini for prominent actin bundles which resemble the rapid loss of E-cadherin from cell -cell contacts^49,50^ This is crucial as it has been shown that E-cadherins can heterotypically bind to N-cadherins during epithelio–fibroblast contact but that these links are highly transient^51^. As epithelial cells strongly form their cell-substrate bonds, they spread and occupy the entire cell-surface interface causing NIH3T3s to phase separate above. Consequently, fibroblasts are highly dependent on the E-N cadherin heterotypic binding to allow adherence. By inhibiting Rock, the E-cadherin focalizing mechanisms which promotes cell-cell contacts are downregulated, resulting in poor adhesion between cell types. As such, NIH3T3s become highly dependent on their homotypic Ncadherin binding and form larger patterned aggregates.

This result is in accordance with previously observed homophilic type binding which favours thermodynamically stable patterns^4,5,15,48^. Cells present a higher affinity to other cells that share their unique cadherin subfamily configuration. This differential adhesion leads to a system with lower interfacial free energy^4,34,48^ It is the differential adhesive strength of these binding affinities that we hypothesize creates a two layered phase separation.

To test whether this two-dimensional patterning phenomenon could result from a minimization of interfacial free energies between binding affinities, we developed a kinetic Monte Carlo simulation. Binding parameters were chosen by translating the homotypic, heterotypic and substrate adhesion forces into relative binding energies. MDCKs’ cell-cell and cell-substrate binding are significantly stronger than the transient bonds that NIH3T3s transiently break in order to migrate^51^. Based on this, we have simulated a co-cultured system whereby cellular movement is solely driven by differential binding affinities. The result is a two layered system in which there is a distinct partition between cell types (Fig 3). Cell type 1 (MDCK) are dispersed and remain adjacently adhered to the substrate whilst cell type 2 (NIH3T3) form aggregates layered above. The simulated cell patterning is similar to what the experimental results showed, which suggests that differential adhesion may be sufficient to dictate cellular organization patterning.

It is not surprising that complex cellular organizational processes such as embryogenesis and tissue repair are governed by mechanisms that exceed any single model of migration. The results shown here however suggest that in cocultured systems, cell-cell and cell-substrate binding affinities are highly influential in dictating cell morphogenesis. We hypothesize that there is a constant interplay between physical forces and biological signalling cues within the microenvironment. Together, they inform the cell’s sensory perception of its surroundings and dictate cellular behaviour. The future lies in further elucidating the balance of this interplay and to more comprehensively understand the mechanisms by which they function.

## Acknowledgements

S.H. was supported by the Queen Elizabeth II Graduate Scholarship in Science and Technology. A.E.P. gratefully acknowledges generous support from the Canada Research Chairs programme. This work was also supported by the Natural Sciences and Engineering Research Council of Canada (NSERC) Grant No. RGPIN/046434-2013 to G.W.S and RGPIN/355535 to A.E.P. We would also like to acknowledge the University of Ottawa for their generous support.

